# The relationship between ageing and changes in the human blood and brain methylomes

**DOI:** 10.1101/2021.01.26.428239

**Authors:** Patrick Bryant, Arne Elofsson

## Abstract

Changes in DNA methylation have been found to be strongly correlated with age, enabling the creation of “epigenetic clocks”. Previously, studies on the relationship between ageing and DNA methylation have assumed a linear relationship, but no study has shown this always to be the case. We show that the relationships between significant methylation changes and ageing are different in different tissues, and that these changes show variable rates across life. Further, we observe a tendency for saturation as ageing proceeds. We provide a straightforward method of assessing all methylation-age relationships and cluster them according to their relative change rates based on fold change. Less than 13% of the significant markers selected here are used in the most common epigenetic clocks. When the significant markers display largely non-linear relationships, our fold change selection outperforms the most common epigenetic clocks in predicting age. We also show that the saturation observed at older ages explains the earlier observations that the epigenetic clock consistently underestimates the age in older samples. The findings imply that markers selected from linear adjustments or correlations do not represent the most meaningful biological changes.

## Introduction

Ageing is a process experienced by most organisms, but why ageing occurs and how to define it on a biological level remains elusive. Epigenetic changes, such as alterations in DNA methylation, histone modifications and changes in chromatin states have been found to be related to ageing in animal models [1], suggesting an epigenetic role for the regulation of lifespan [2].

In particular, the role of DNA methylation on CpG sites has been used to relate chronological age (CA) to DNA methylation age (DNAmAge), deemed the “epigenetic clock” [3]. The original clock, by Steve Horvath, uses a linear combination of 353 CpG methylation sites selected by elastic net regression [4] and can predict CA with a correlation of 0.96 and an average error of 3.6 years, using 51 healthy tissues. Another model by Hannum [5], using only blood, uses the same principle to obtain a correlation of 0.96 and an accuracy of 3.9 years.

Non-linear models have been used to improve the DNAmAge predictions resulting in higher correlations between DNAmAge and CA [6],[7]. These models were built using preselected methylation markers, using elastic net regression and analysis of Pearson correlations, thereby favouring selecting probes based mainly on only linear relationships between CA and DNA methylation levels.

However, there have been reports of “systematic underestimation of the epigenetic clock and age acceleration in older subjects” [8], suggesting that the relationship between methylation change and CA is non-linear. However, it may have a sizable linear regime since the non-linear effect is most prominent in older individuals. The underestimation was observed in all examined tissues, but most apparent in the Cerebellum.

Methylation levels at only a few CpG sites, used in epigenetic clocks, can be sufficient for predicting ageing, and accompanied death risk to a certain extent [3,5,9,10]. Attempts to analyse the real effect on gene expression of these experimentally, found hypomethylation being associated with ageing [11]. Longitudinal studies have further attempted to validate methylation markers by looking at marker consistency across measurements and their relation to ageing and disease [12,13].

The studies mentioned above assume a linear relationship between ageing and important methylation markers by either correcting for ageing or correlating age with methylation change in a linear combination. Even though a linear feature combination allows for a nonlinear prediction, selection of features using linear methods favours constant, linear relationships. However, there is no proof that this assumption is valid, i.e. that the most important relationships between ageing and methylation is linear. Therefore, we here compare the relationship between methylation markers and ageing without making assumptions on the relationship between methylation and ageing.

## Methods

### QC and grouping

We utilise Illumina Infinium 450k Human DNA methylation profiles from a set of 656 blood samples from the ArrayExpress database (E-GEOD-40279, https://www.ebi.ac.uk/arrayexpress/experiments/E-GEOD-40279/) [5], 428 frontal cortex and 402 cerebellum samples (E-GEOD-36194, https://www.ebi.ac.uk/arrayexpress/experiments/E-GEOD-36194/ and E-GEOD-15745, https://www.ebi.ac.uk/arrayexpress/experiments/E-GEOD-15745/) [14]. The samples span a vast range of ages (19 to 101 years for blood, and 0 to 102 years for the frontal cortex and cerebellum, Figure 1) and have balanced sex distributions. The methylation normalisation and quality control were used as described in the original publications (see E-GEOD accessions).

**Figure 1.**
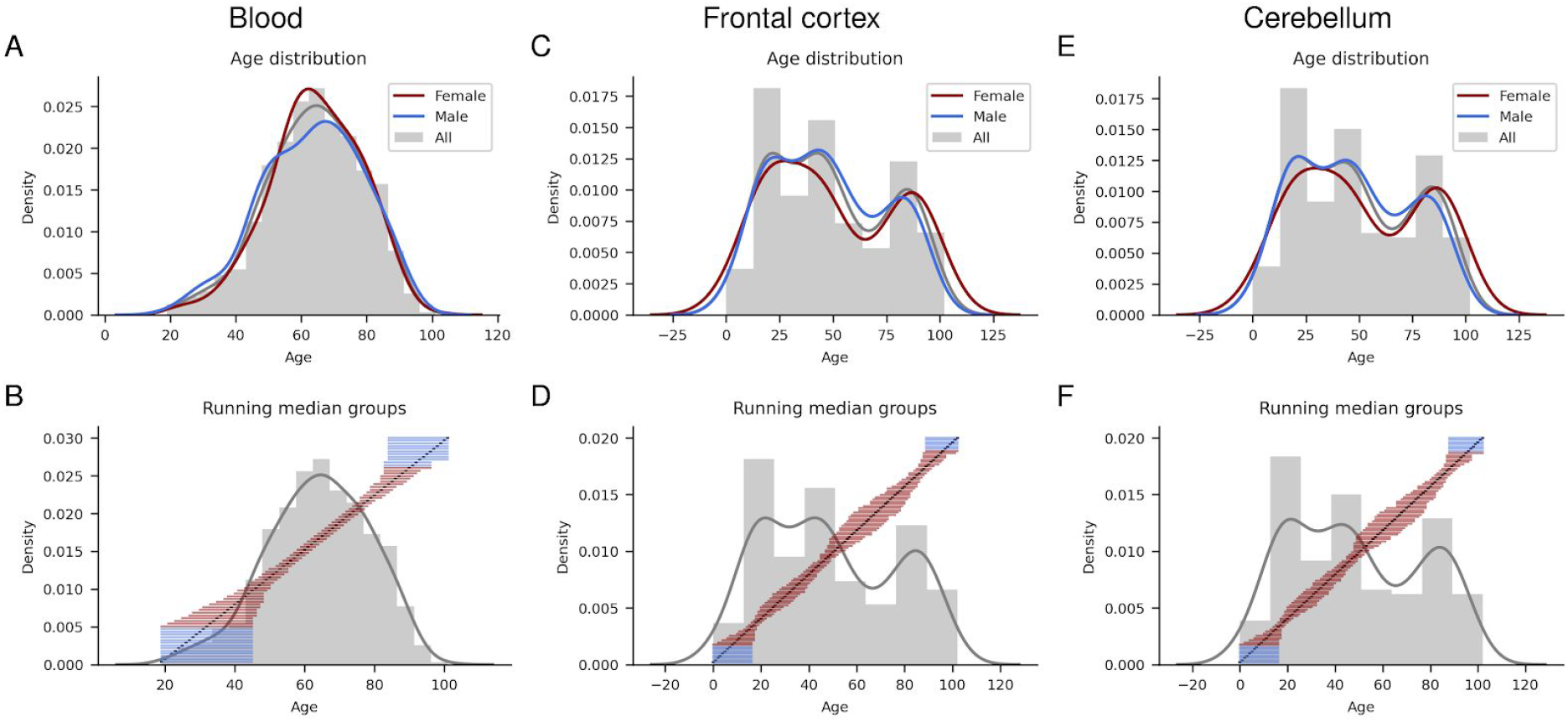
**A, C, E** Age distribution of all samples and divided by sex for blood, frontal cortex and cerebellum respectively. The ages of the samples range from 19 to 101 years for blood and from 0 to 102 years for Frontal cortex and Cerebellum. **B, D, F** The inclusion limits for the running median groupings using the closest 10 % of samples for each year displayed as horizontal blue and red lines. The red lines show the groups used in the analysis, while blue show those that were excluded. The age histogram in grey and the age being grouped marked in black, blood, frontal cortex and cerebellum respectively.

Further quality control was performed by removing markers with over 10 missing beta values (40110 out of 473034 from the blood samples and 0 out of 27476 from the frontal cortex and cerebellum samples). We then calculated the mutual information [15] for the beta-value distribution of each sample by comparing with the mean beta value distribution. We removed samples with a mutual information score of less than three standard deviations away from the mean (eight from blood, two from frontal cortex and five from Cerebellum, see Figure S1).

### Investigation of methylation-age relationships

To investigate not only linear relationships between marker methylation change and age. We also create a running median of the methylation beta values for each probe. The median is less sensitive to outliers compared to an average. To avoid grouping effects, we select 10% of the closest samples by age for each age with steps of one year from the lowest to the highest age. For example, the starting age for blood, 19, only has one sample, why samples up to the age of 45 are included in the median calculation. By including samples selected by age distribution, it is ensured that each calculation has equal sampling.

To avoid edge effects that may arise due to low sample representations at high and low ages, where the samples are fewer, only the groups whose midpoints are close to the ages being represented were used in the analysis (ages 32-90 years for blood, 8-96 for frontal cortex and 8-95 for cerebellum). From these running medians, the maximum fold change (FC) for each marker is assessed, by calculating the difference between the minimum and maximum methylation medians.

A two-sided t-test was made by comparing the samples used to calculate the maximum FC observed for each marker. The markers with a false discovery rate (FDR) below 0.05 (according to the Benjamini-Hochberg procedure [16] in statsmodels[17]) and having an FC higher than two were selected. Selected markers were analysed both individually and by their relationships to genes [18], [19]. To ensure that the median values are stable, we analysed the standard deviation for each interval used. We compute the relative standard deviation by dividing the standard deviation with the median in each interval. If the average relative standard deviation is above 0.5, we consider the spread to be too large and do not include this methylation marker in the analysis.

To analyze the tendency for marker beta values being linearly related to ageing, the gradients of the running medians were calculated. The beta-values for the gradients were normalised with the highest beta value for each marker.To obtain the same relative change for each marker and thus make them comparable. The gradients were thereafter clustered using k-means clustering [20]. To select the number of clusters, t-SNE embeddings were computed from the gradients [21]. The number of clusters was then chosen after analysis of the first two components of these embeddings, ensuring sound separation of the resulting two-dimensional points. The gradients are noisy and were for visualisation purposes, therefore, smoothed using a Savitzky-Golay filter (with window length 21 and polynomial order 2) [22].

### Gene Ontology enrichment

To analyze the biological importance of the genes related to the selected markers, Gene Ontology (GO) enrichment was performed using PANTHER, version 14 [23]. The enrichment was performed for each cluster in each tissue, to analyze potential differences and similarities between markers with different age relationships.

### Statistical significance for the overlap of two sets of markers

To calculate the probability of the selected markers overlapping with those from the Hannum and Horvath epigenetic clocks, one has to calculate the probability of selecting first the markers from each selection and then consider the probability that a certain number of these match. This is done by consulting the Hypergeometric distribution or a normal approximation of this since the exact hypergeometric probability is difficult to calculate (due to the large products for large factorials and the problem with representing these in computers).

x = number of markers in common between two groups.

n = number of markers in group 1.

D = number of markers in group 2.

N = total number of markers

The normal approximation is used when:

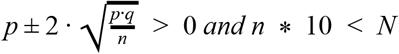

where p = D / N and q = 1 - p.

The normal approximation is:

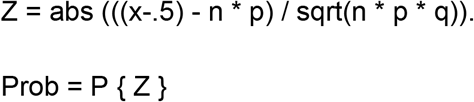

where Z is a standard normal variation from N(0,1).

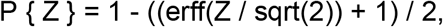

where erff is the error function, see http://nemates.org/MA/progs/overlap_stats.html for code, calculations and further details.

### Comparing age predictions from linear and fold change selection

To ensure the presented fold change selection results in more informative markers, we build a machine learning model in the form of a random forest regressor with scikit-learn [24] using the selected markers. We compare the accuracy from our model with that of predictions using both the Hannum model for blood (blood only) and the Horvath model (all tissues) for the Cerebellum and Frontal cortex in a 5-fold CV. To reduce training bias and display the strength of the selected markers, we donät optimize any model parameters. We perform training on 80 % of the data and validate the remaining 20 % using the default parameters in scikit-learn.

## Results

### The relationship between significant methylation changes and ageing

There are 120 out of 432924 methylation markers for blood, 514 out of 27476 for the frontal cortex and 152 out of 27476 for the cerebellum, which running medians display a fold change (FC) above 2 that are statistically significant on false discovery rate (FDR) 0.05. The p-value distributions are sound and suggest a well-calibrated experiment (Figure 2). All significant markers for blood, frontal cortex and the cerebellum are divided into 2, 1 and 4 clusters respectively (Figures 3, 4 and 5).

**Figure 2.**
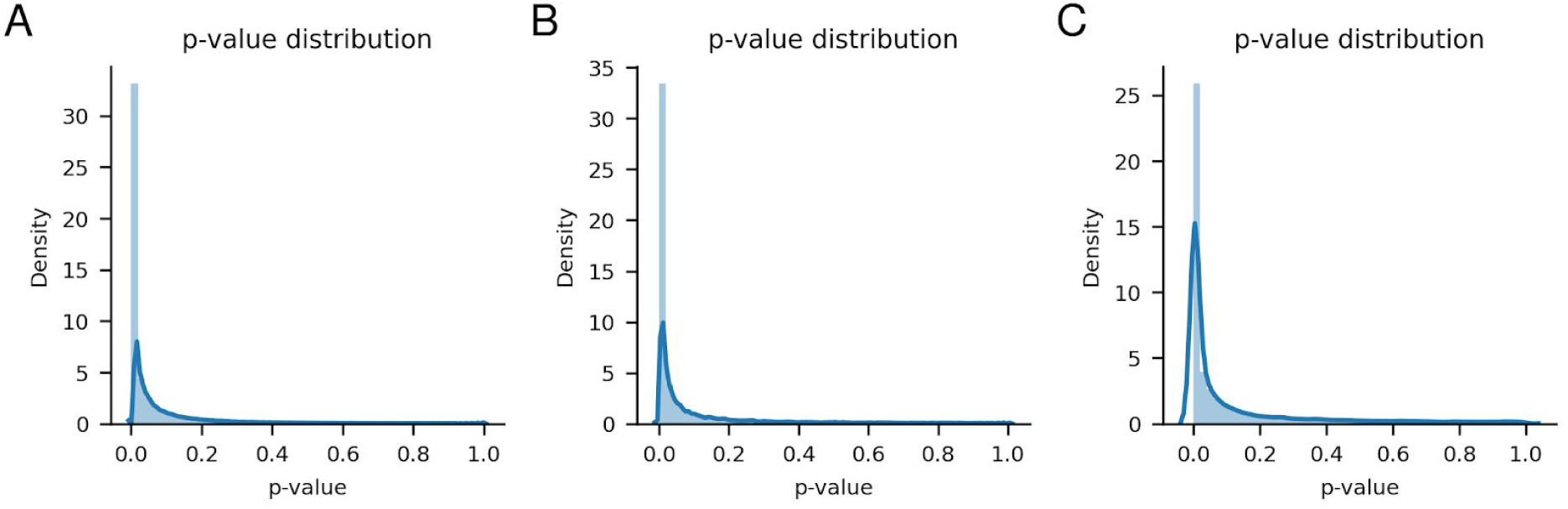
**A, B, C** p-value distributions for blood, frontal cortex and the cerebellum respectively.

**Figure 3.**
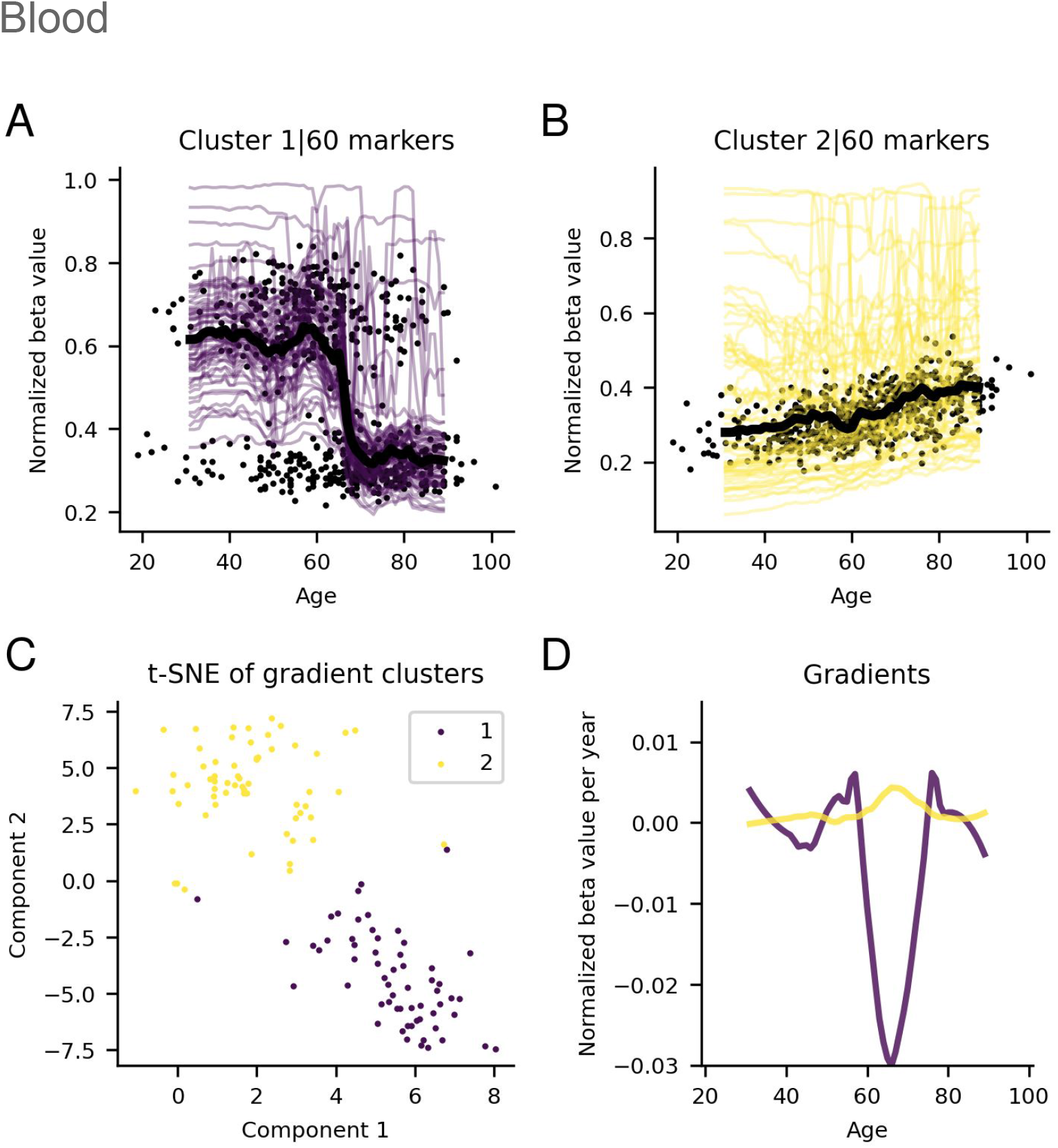
**A, B** Significant running medians for marker clusters from the blood. The black points represent the median value for each sample and the black lines the running median for each cluster. The beta values have been normalized with the highest beta value for each marker. **C** Visualization of the t-SNE cluster embeddings on two components. **D** Gradients of running medians against age, smoothed medians using a Savitzky-Golay filter [22] with window length 21 and polynomial order 2.

**Figure 4.**
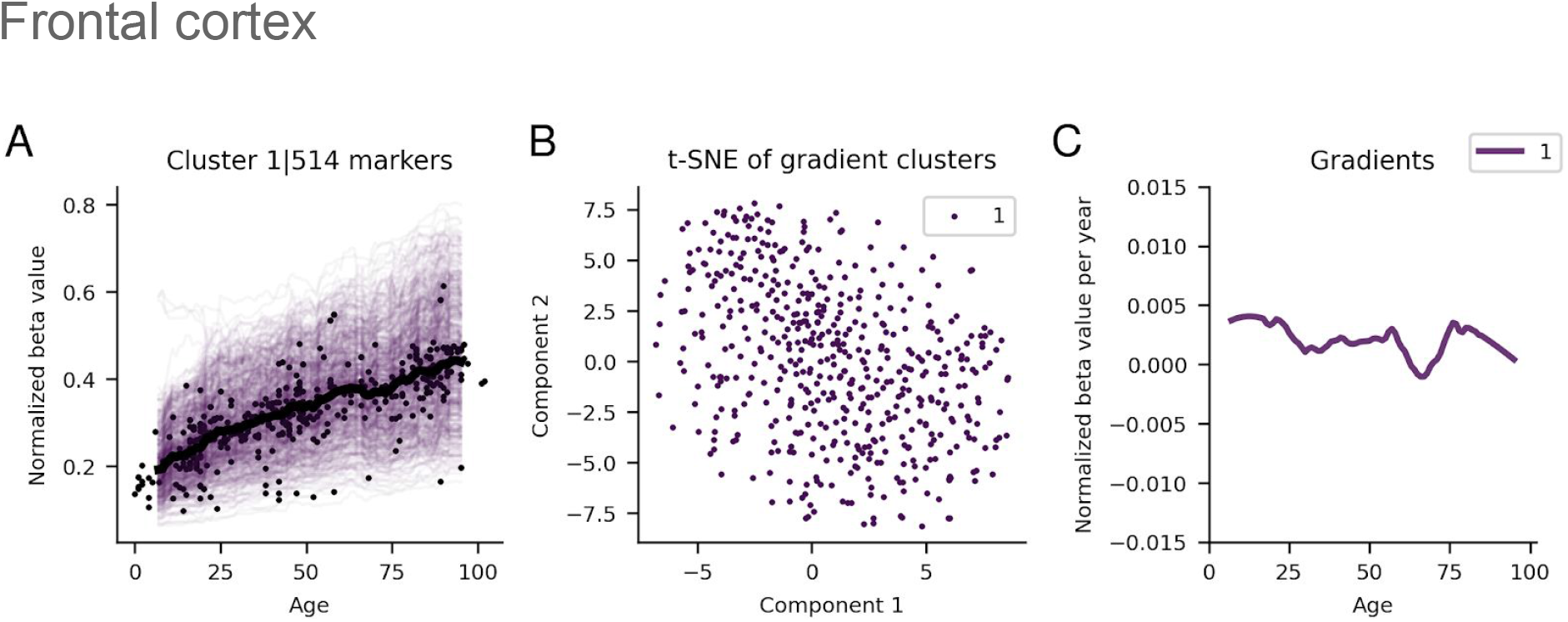
**A)** Significant running medians for the marker clusters from the frontal cortex. The black points represent the median value for each sample and the black lines the running median for each cluster. The beta values have been normalised with the highest beta value for each marker. **B)** Visualization of the t-SNE cluster embeddings on two components. **C** Gradients of running medians against age, smoothed medians using a Savitzky-Golay filter [22] with window length 21 and polynomial order 2.

**Figure 5.**
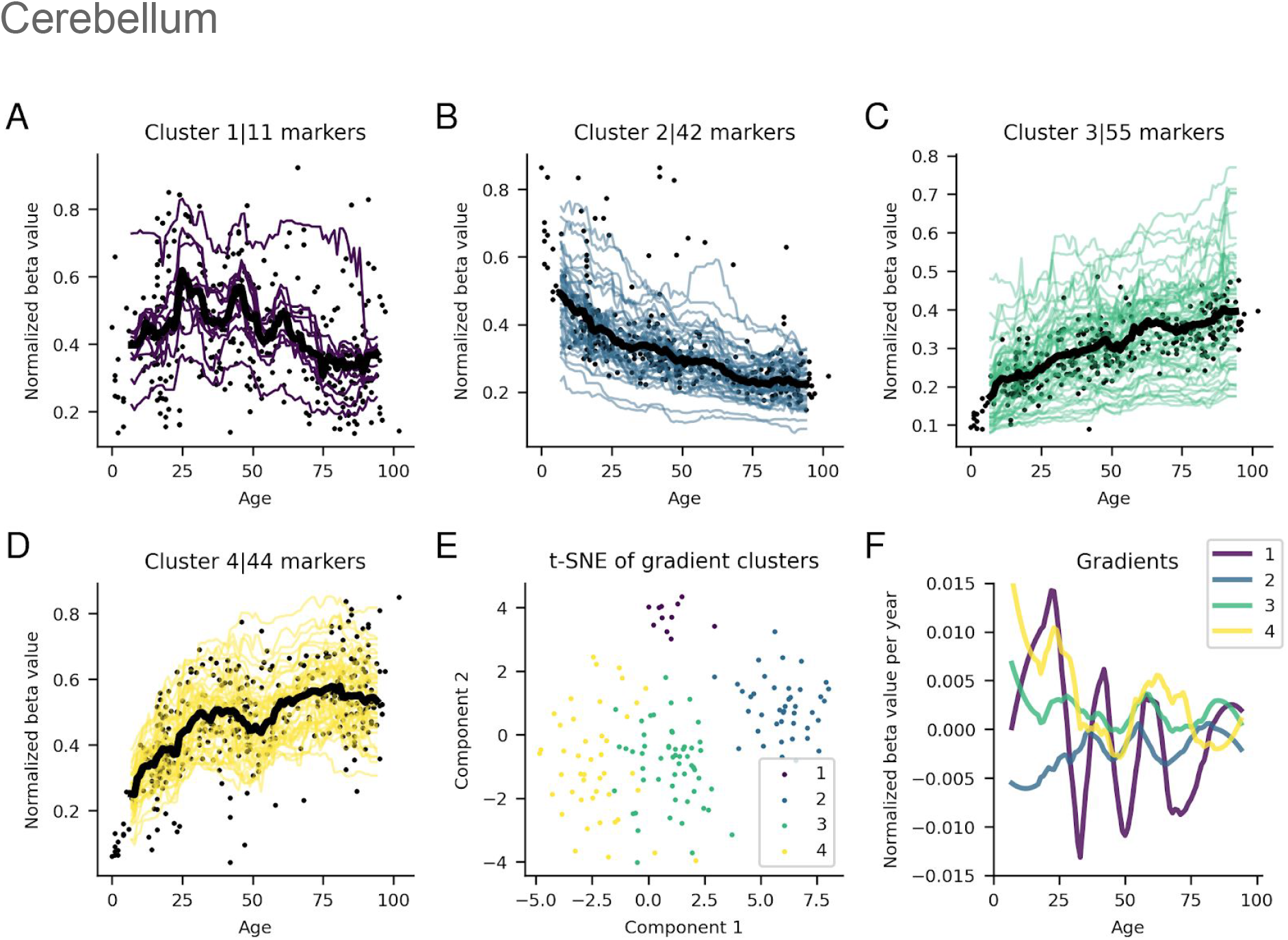
**A-D** Running medians for significant marker clusters from the cerebellum. The black points represent the median value for each sample and the black lines the running median for each cluster. The beta values have been normalized with the highest beta value for each marker. **E)** Visualization of the t-SNE cluster embeddings on two components. **F)** Gradients of running medians against age, smoothed medians using a Savitzky-Golay filter [22] with window length 21 and polynomial order 2.

One of the clusters found in blood displays negative correlation (cluster 1, 60 markers) and the other positive (cluster 2, 60 markers) correlation with age (Figure 3). These clusters show clear separation from each other, and contain only a few outliers. For the positive relationship in blood, the methylation rate is steady from age 32 to age 60, where it rapidly accelerates to age 70, to slow down and flatten out for older ages. The negative relationship in the blood is a more extreme inverse of the positive, steady states before and after a large drop around age 60 are observed. The beta values for both clusters are quite low (0.2-0.1), with outliers of higher values (Figure S2 A and B).

In the frontal cortex (Figure 4), only one, central cluster with 514 markers was found. This cluster only displays significantly positive relationships with ageing. The gradients for this cluster are very stable throughout all ages (8-96 years). At age 8, the gradients are positive, with a median change of about 0.4% of the maximal observed beta value per year. As ageing proceeds, the gradients decline towards zero. The number of points rapidly declines, and the spread increases in this area. The beta values range from close to 0 to 0.1-0.2 (Figure S2 C).

For the cerebellum, four distinct clusters were found, with varying degree of separation (Figure 5). Cluster 1, which is clearly separated from the others, contains 11 markers and displays an up-down relationship, with increasing methylation up to age 25, to rapid decrease and flattening out at around age 75. Most beta values in cluster 1 range from 0.1 to 0.5 (Figure S2 D). The gradients there are very noisy though, likely due to the mere 11 markers. Clusters 3 and 4 are adjacent to each other and have 55 and 44 markers respectively. These clusters show similar relationships with ageing, accelerating methylation up to age 30 and flattening out towards older ages. Cluster 4 has a much more rapid acceleration though, resulting in a higher steady-state normalized beta value compared with cluster 3 (beta values of 0-0.2 and 0.1-0.6 respectively, Figure S2 F and G). Cluster 2, separated due to its negative relationship with ageing, has 42 markers and displays the inverse relationship of cluster 2. The beta values in this cluster range from 0.1-0.6 (Figure S2 E).

### Comparing marker selections from direct age correlations, epigenetic clocks and running medians

Out of the 120 significant methylation markers for blood, 514 for the frontal cortex and 152 for cerebellum 112, 510 and 147 markers were significant on FDR 0.05 when analyzing the Pearson correlation coefficient between age and marker values. The correlation analysis deemed 190238 (44%), 10235 (37%) and 6022 (22%) unique markers significant on FDR 0.05 for blood, frontal cortex and cerebellum respectively. When selecting markers with significant age correlations (FDR<0.05), one obtains a bimodal distribution of Pearson correlations (Figure 6). Another distribution is obtained from the running median selection, showing that even poorly correlated markers can portray significant methylation changes for the blood and the cerebellum, although the frontal cortex selection shows almost only positively correlating markers.

**Figure 6.**
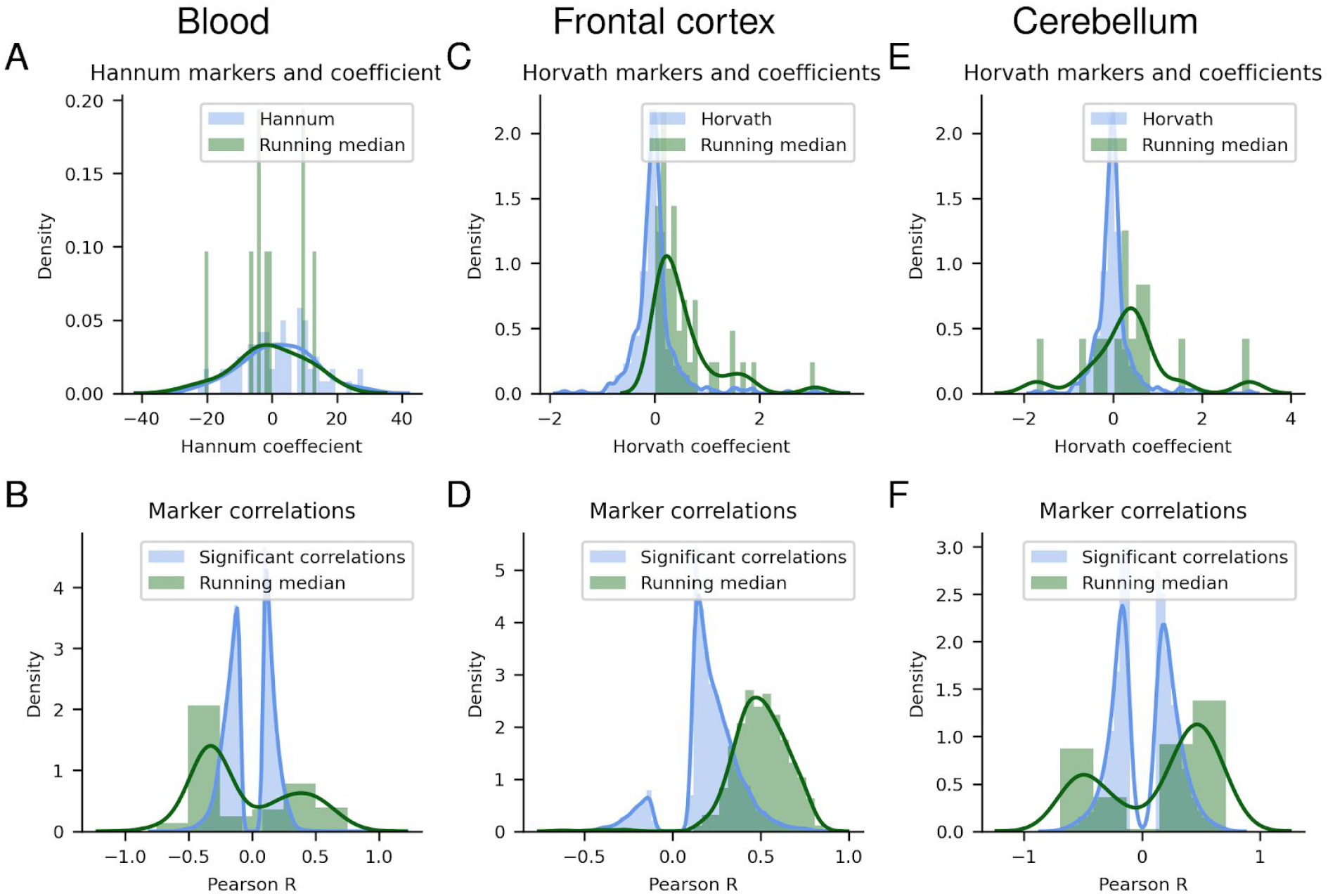
**A, C, E** The coefficients for the linear marker combination in the Hannum (71 markers) and Horvath (353 markers) clocks and the overlapping markers selected from the running medians in Blood, Frontal Cortex and Cerebellum respectively. **B, D, F**) The distribution of the Pearson correlation coefficients from the markers with significant age correlations and those selected from the running medians. Blood, Frontal Cortex and Cerebellum, respectively.

Of the 120 significant blood markers, only 9 overlaps with the 71 markers that are used in the Hannum epigenetic clock [5], all belonging to cluster 2 (Figure 3). Of the 514 and 152 significant markers in the frontal cortex and cerebellum, 41 and 15 overlap with the 353 markers that are used in the Horvath epigenetic clock [3], respectively. All 41 in the frontal cortex belong to the only cluster (Figure 4), while of the 15 in the cerebellum 11 belong to cluster 3 and 4 to cluster 2 (Figure 5). The overlapping markers have both positive and negative coefficients for the cerebellum and blood, while the frontal cortex has coefficients of 0 and above (Figure 6). The probability of selecting the significant markers randomly and that the observed overlap with the epigenetic clock markers would be found is p < 5.207e-22, p < 9.290e-21 and p < 1.301e-09 for blood, frontal cortex and cerebellum respectively, see methods section.

### Comparing the prediction accuracy between epigenetic clocks and running median selections

The results from fitting a random forest regressor to the running median selections display a reduction in error compared to the Horvath clock for the cerebellum (6.65 vs 9.48 years), but a slightly higher error for the frontal cortex (6.79 vs 5.35 years), see Figure 7 and Table 1. For blood the error is 5.52 years for the Hannum clock vs 5.03 years for the running median selection, even though the Hannum clock has been trained on the same dataset [5]. The Pearson correlation coefficients are lower for the random forest models in blood and the frontal cortex, but higher for cerebellum. The previously reported underpredictions for the cerebellum [8] manifest here as well.

**Figure 7.**
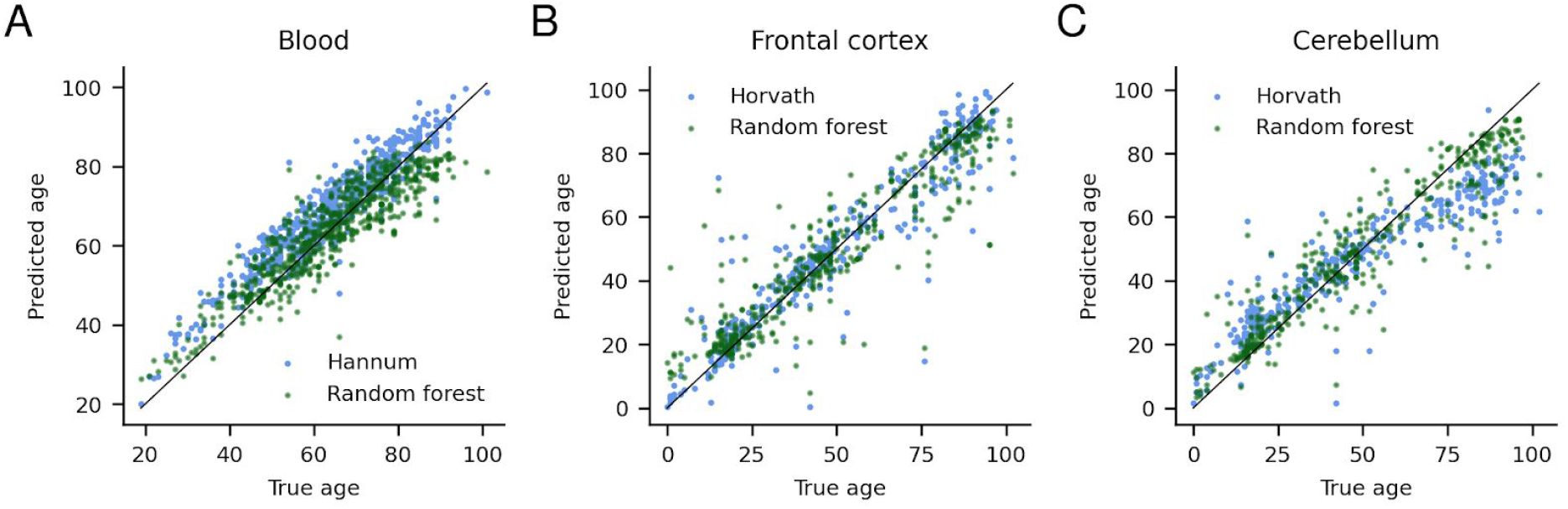
Results from fitting a random forest regressor to the blood **(A)**, frontal cortex **(B)** and cerebellum **(C)** markers and comparing with the Hannum (blood) and Horvath (frontal cortex and cerebellum) results. One can see that the results are substantially improved using the running median selection method for the cerebellum, but not for the frontal cortex or blood.

**Table 1.** Average errors and Pearson correlations for the Hannum, Horvath and random forest models fitted to each tissue type.

| Model | Error | PCC |
| --- | --- | --- |
| Blood Hannum | 5.52 | 0.95 |
| Blood Random Forest | 5.03 +/- 0.28 | 0.90 +/- 0.01 |
| Frontal cortex Horvath | 5.35 | 0.95 |
| Frontal cortex Random Forest | 6.79 +/- 0.61 | 0.92 +/- 0.02 |
| Cerebellum Horvath | 9.48 | 0.93 |
| Cerebellum Random Forest | 6.65 +/- 0.72 | 0.94 +/- 0.01 |

### Genes regulated by significant methylation markers and GO enrichment

Since several markers may regulate the same gene, the markers were grouped by gene and further analysed. The relationship of all genes with at least two significant methylation markers was assessed to investigate potential differences in methylation change for these markers. There are 99, 438 and 140 unique genes of which 5, 32 and 5 are regulated by at least two significant markers for blood, frontal cortex and cerebellum, respectively. The 5, 32 and 5 genes are regulated by 11, 65 and 10 markers, respectively. All of these show positive relationships with age (Figures S3, S4 and S5), belonging to clusters 2, 1 and 4 for blood, frontal cortex and cerebellum respectively.

GO enrichment of the unique genes suggests what processes the markers regulate (Figures S6, S7 and S8). The two marker clusters for blood are mostly related to metabolic and cellular processes, as well as biological regulation with similar frequencies. This is true for the Hannum selection as well, although both cluster 1 and the Hannum selection show relationships with more different processes than cluster 2. The frontal cortex enrichment (only one cluster) shows a wide variety of processes, with a very large overlap with the Horvath marker GO terms. The four different cerebellum clusters show a wide variety of GO terms, although metabolic processes, cellular processes and localization are important in all. Clusters 2 and 3 seem to have most overlap with the Horvath terms. We note that these are the clusters with the most linear relationships. All clusters show important differences, supporting the validity of their division.

## Discussion

An issue with investigating marker correlation with age is not evaluating the fold change (FC), only the correlation itself. A very high number of markers are significant on FDR 0.05 from the Pearson correlation analysis. However, a marker that changes only 1 % in total, but does so with 0.01 % a year from year 0-100 will have a Pearson correlation coefficient of 1.0 with age. When evaluating the FC, larger relative methylome changes are assessed, thereby capturing changes that likely have a higher impact on gene expression.

The number of individuals for each age is not uniform, which is usually the case in clinical studies (see Figure 6). We provide a solution to this problem by selecting 10% of the closest samples in age for each year and thus creating equal sample representations for each age. Further, by comparing the maximal FC for the running median across all ages, we ensure all significant changes throughout ageing are assessed. The t-test of the significance between the samples used to construct the median points related to the FCs ensures statistical soundness.

To investigate all possible significant relationships between ageing and methylation change, k-means clustering was performed on the normalized gradients. The number of clusters was evaluated through t-SNE, ensuring sufficient separation between gradient clusters. No cluster in any tissue displays methylation-age relationships equal to another. This suggests age-related methylation is highly tissue-specific, as suggested from studies of tissue-specific age predictors [7].

What is common in all methylation-age relationships is a tendency for saturation as ageing proceeds. This explains why there is a consistent underprediction of the age in older samples for the epigenetic clocks that use linear combinations of methylation change [8]. This tendency should not be due to underrepresentation of older samples, as the equal age representations and control for edge effects (see methods) prevent such issues. Further, no relationship is found to be strictly linear, although the one found in the frontal cortex is quite close. Analysis of the gradients, however, provides a more clear picture of how the medians change, something that is difficult to capture when analyzing only the direct relationships.

All tissues contain clusters which have linear regimes. This is easy to identify for the frontal cortex and cerebellum. For blood, however, cluster 1 has only a very narrow window which could be considered semi-linear, the region of rapid methylation increases between ages 60 and 80 (Figure 2). This provides an answer to why the epigenetic clocks and age adjustments in previous studies [3,5,9–13] work to a certain extent, although markers with non-significant changes have mostly been used in these.

Blood is the tissue with the most protrusive methylation relationships. Why the methylation levels change rapidly at age 60 is unclear. An explanation may be a downregulation of immune responses at older ages [25], as DNA from blood is mostly from Leukocytes [26]. The running medians constituting the blood clusters have substantial spread as well. This creates noise for the total cluster medians (black lines in Figures 3,4 and 5), which may impact on the observed relationships. The Savitzky-Golay filter applied to the gradients should be able to counteract such noise, capturing significant signals.

The frontal cortex only displays hypermethylation, while both blood and the cerebellum display both hypo- and hypermethylation. Since hypomethylation is related to increased gene expression [27,28], the continuous hypermethylation in the frontal cortex throughout ageing suggests gene expression is continuously downregulated. This corresponds well with previous findings in the prefrontal cortex, where DNA methylation changes have been found to be fast during the prenatal period, and later slow down and do so continuously with ageing [29].

The beta values were normalised to be able to compare relative change in methylation and thus obtain comparable gradients. In Figure S2, unnormalised relationships are displayed. The beta values for blood and the frontal cortex are very close for the majority of the running medians, but low, only amounting to maximal 20% methylation. Those for the cerebellum, on the other hand, are both larger and show greater variability, most ranging from 0.1-0.5. This suggests that the gene expression in the cerebellum may be more drastically altered during ageing, compared to the more fine-tuned changes observed in the other tissues.

When comparing the significant markers with those of the Hannum (only blood) [5] and Horvath (multi-tissue) [3] epigenetic clocks, there is significant overlap (p < 5.2 · 10^-22^, p < 9.3 · 10^-21^ and p < 1.3 · 10^-9^ for blood, frontal cortex and cerebellum respectively). Despite the significant overlap, less than 15% of the significant markers are overlapping (13, 8 and 10% for blood, frontal cortex and cerebellum respectively), suggesting that 87-92% of the significant changes are missed by linear analysis. The argument for selecting markers that correlate highly with age is thus still without support, something highlighted in the non-linear findings here.

That the random forest prediction errors are better in the cerebellum compared to the Horvath clock, while the frontal cortex shows similar performance, can be explained by the fact that the markers selected in the frontal cortex have a more linear relationship with age. This is also true for blood (cluster 2 representing half of the markers show a linear relationship), which shows only a slight improvement with the random forest model. When this is not the case, as in the cerebellum, the fold change selection method presented here clearly outperforms the linear selection. It should be noted that the markers selected here were also not chosen for the objective of predicting age, which is the case in the epigenetic clocks. This suggests that selecting markers based on fold change enables capturing significant biological age relationships, while linear selection only manages when the age relationships happen to be largely linear.

The GO enrichment shows sound annotations for all tissues and epigenetic clock comparisons. Interestingly, different clusters display GO terms related to different biological processes, within the same tissue. The overlap with the epigenetic clocks is greatest for the clusters with more linear relationships. Without evaluating the effect size of the methylation change, it is difficult to know if a marker will have much impact on gene expression or development.

## Conclusions

The relationship between significant DNA methylation changes and ageing varies between tissues and life periods. The variability in the relationship between methylation and age, especially the saturation observed at older ages, might provide important insights. The regions of semi-linearity explain why epigenetic clocks and age adjustments that assume linear relationships produce results that can be interpreted to be successful. The saturation explains why ageing is underpredicted in older samples. When analyzing the relationship between methylation levels and age, therefore, one has to evaluate not only linear relationships between DNA methylation and age but also non-linear relationships. We show that when significant markers have non-linear relationships with age, the fold change selection outperforms the epigenetic clocks in predicting age. This is true even though the objective of age prediction was not sought after and no parameter optimization was performed for the models built with these markers. We further argue for the importance of fold change for the selection of important methylation markers, because markers with low fold change are unlikely to have a substantial impact on gene expression. Therefore, epigenetic studies and clocks that have used markers selected from only linear adjustments or correlations are unlikely to represent meaningful biological changes. Here, we provide a new way to investigate all possible methylation-age relationships and group them according to their relative change rates.

## Declarations

### Ethics approval and consent to participate

The data is from previous studies, available from ArrayExpress. We refer to the accession numbers E-GEOD-40279, E-GEOD-36194 and E-GEOD-15745.

### Availability of data and materials

The datasets used, E-GEOD-40279

(https://www.ebi.ac.uk/arrayexpress/experiments/E-GEOD-40279/), E-G EOD-36194 (https://www.ebi.ac.uk/arrayexpress/experiments/E-GEOD-36194/) and E-GEOD-15745 (https://www.ebi.ac.uk/arrayexpress/experiments/E-GEOD-15745/), are available at the ArrayExpress database.

The code is freely available under the GPLv3 license: https://github.com/patrickbryant1/ageing_of_blood_and_brain

### Authors’ contributions

Patrick Bryant conceived and designed the experiments, performed the experiments, analysed the data, prepared figures and/or tables, authored or reviewed drafts of the paper, and approved the final draft.

Arne Elofsson analysed the data, authored or reviewed drafts of the paper, and approved the final draft.

## Acknowledgements

We want to thank everyone involved in constructing the datasets E-GEOD-40279, E-GEOD-36194 and E-GEOD-15745 for submitting their data to ArrayExpress as well as everyone involved in developing this magnificent data repository. We also acknowledge the creation of the original epigenetic clocks by Hannum and Horvath.

