## Supplementary material for "The relationship between ageing and changes in the human blood and brain methylomes"

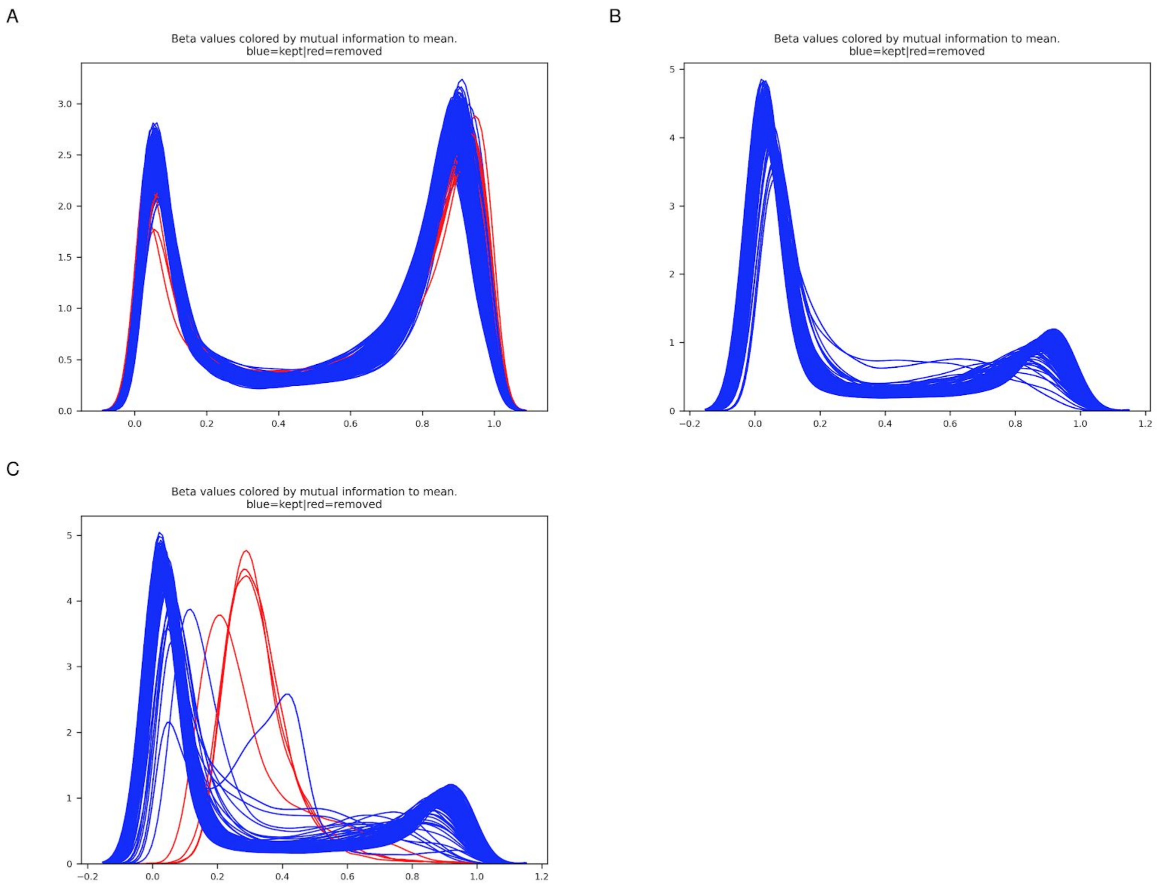


**Figure S1.** Beta value distributions colored by deviation according to mutual information score (see methods). **A** Blood **B** Frontal cortex **C** Cerebellum**.**

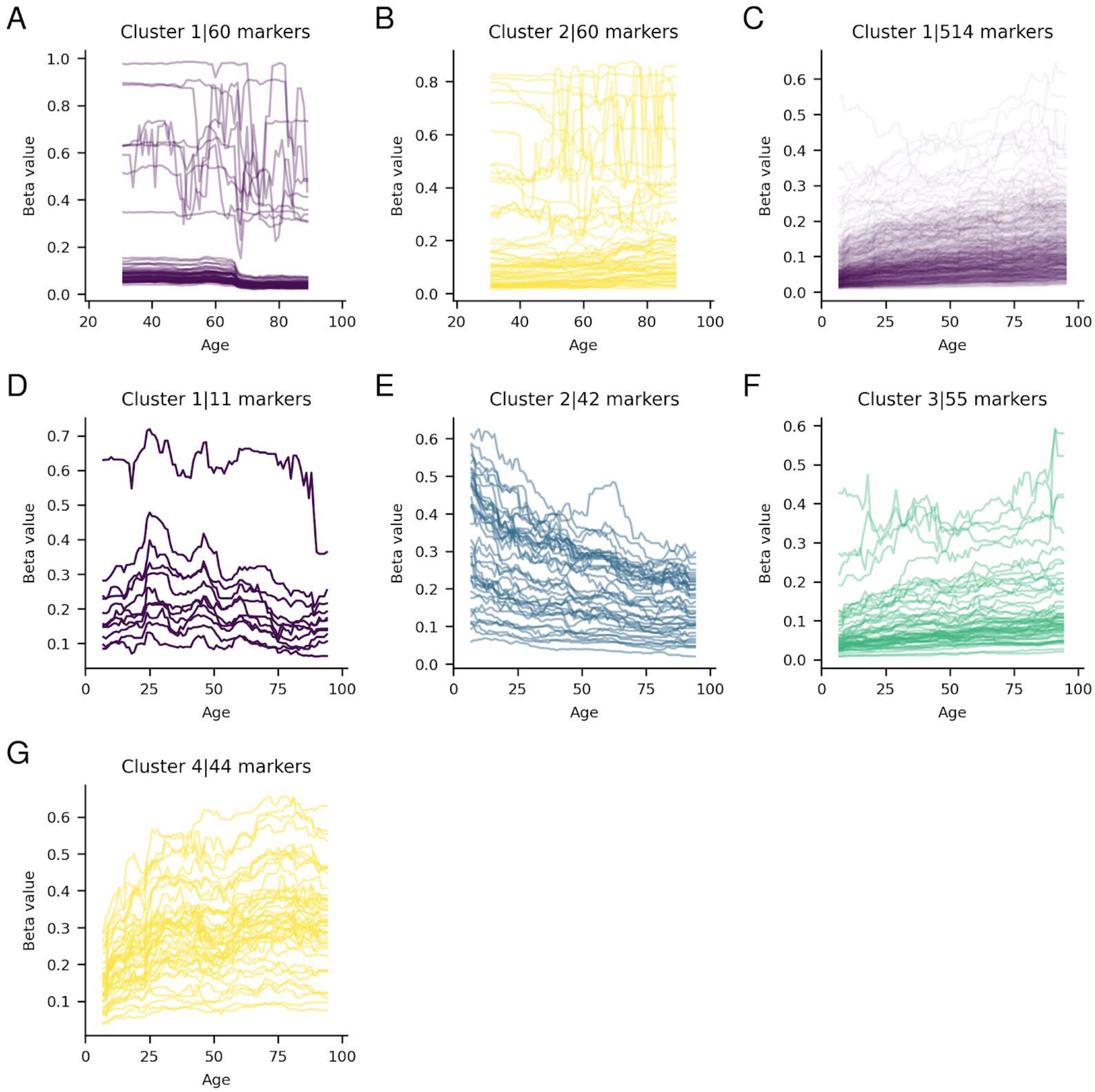


**Figure S2.** Unnormalized significant running medians for marker clusters for blood **(A, B)**, frontal cortex **(C)** and the cerebellum **(D,E,F,G)**.

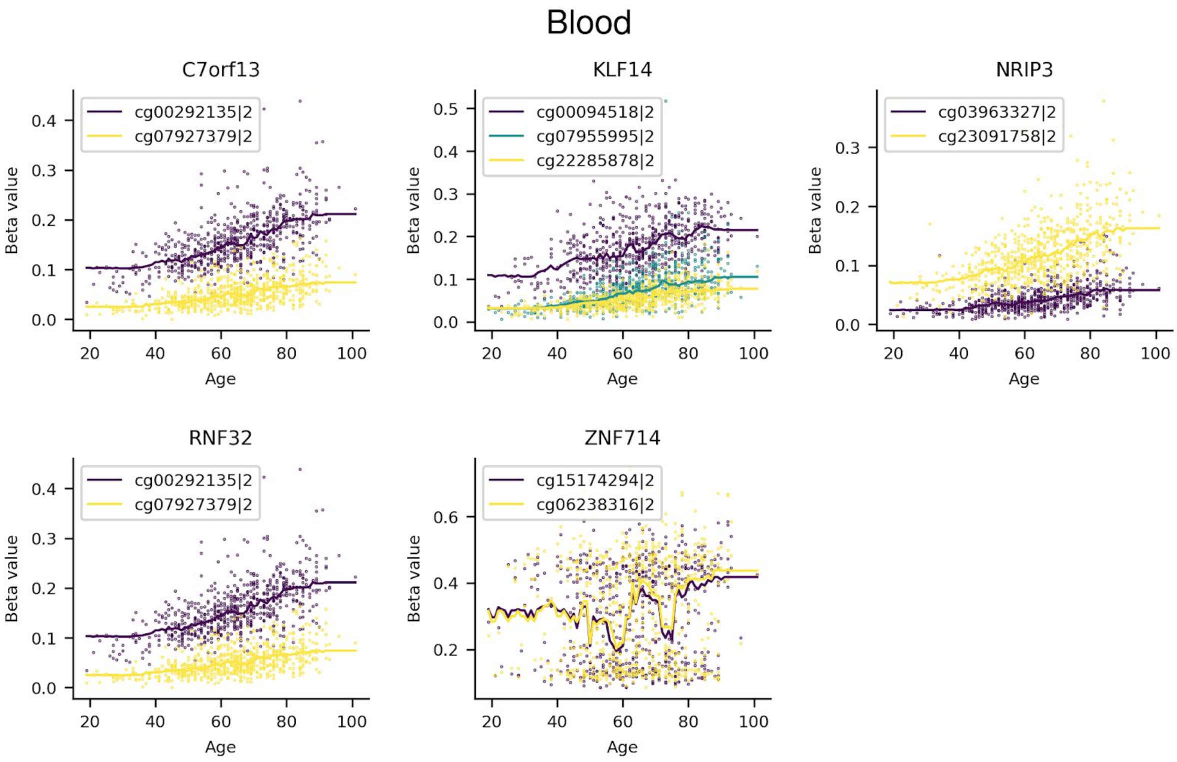


**Figure S3.** Genes for Blood that are regulated by at least 2 significant markers. The marker identifier and the cluster label are separated by “|”. All belong to cluster 2.

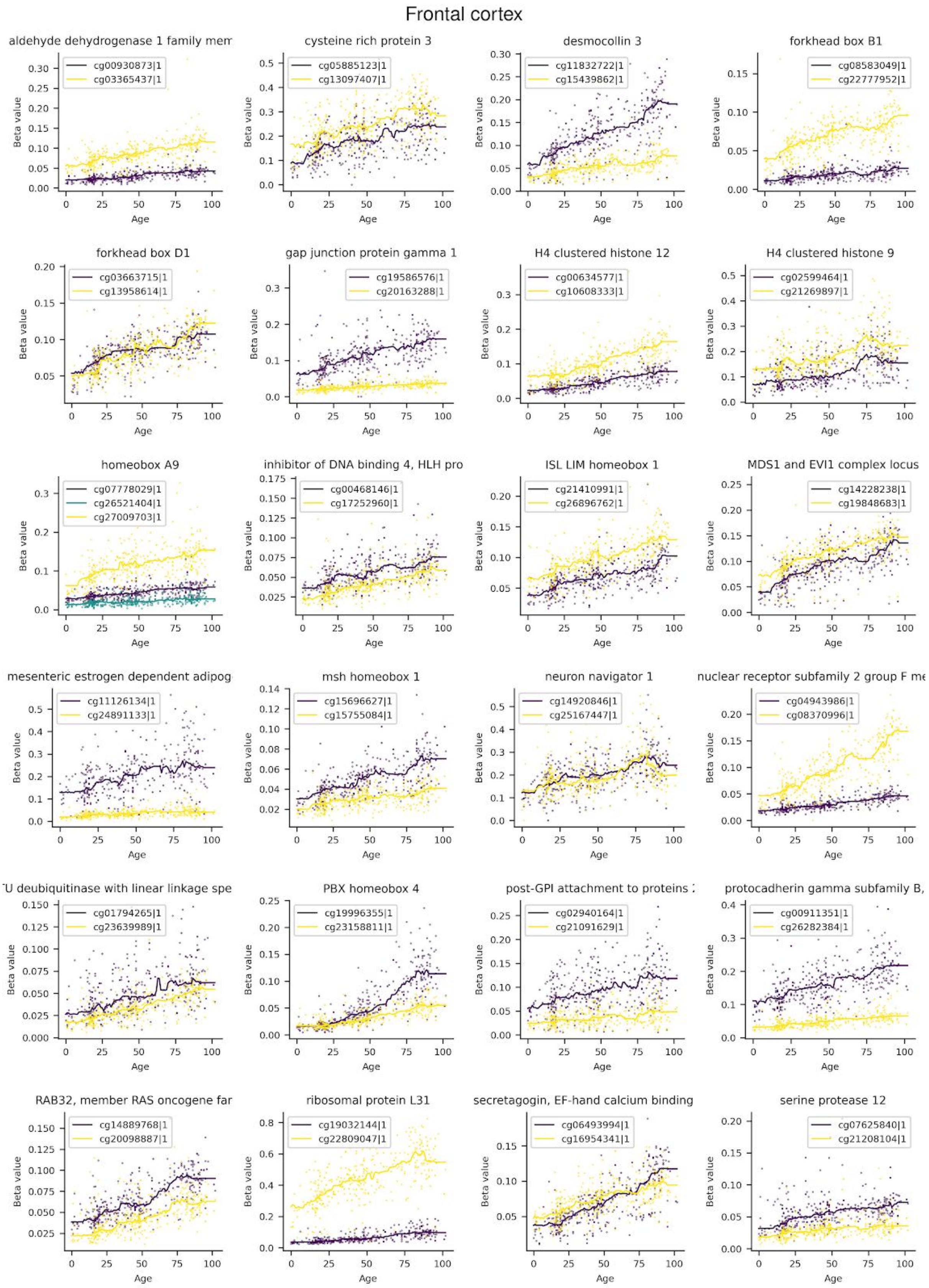
Frontal cortex
aldehyde dehydrogenase 1 family mem
cg00930873|1
cg03365437|1
cysteine rich protein 3
cg05885123|1
cg13097407|1
desmocollin 3
cg11832722|1
cg15439862|1
forkhead box B1
cg08583049|1
cg22777952|1
forkhead box D1
cg03663715|1
cg13958614|1
gap junction protein gamma 1
cg19586576|1
cg20163288|1
H4 clustered histone 12
cg00634577|1
cg10608333|1
H4 clustered histone 9
cg02599464|1
cg21269897|1
homeobox A9
cg07778029|1
cg26521404|1
cg27009703|1
inhibitor of DNA binding 4, HLH pro
cg00468146|1
cg17252960|1
ISL LIM homeobox 1
cg21410991|1
cg26896762|1
MDS1 and EVI1 complex locus
cg14228238|1
cg19848683|1
mesenteric estrogen dependent adipog
cg11126134|1
cg24891133|1
msh homeobox 1
cg15696627|1
cg15755084|1
neuron navigator 1
cg14920846|1
cg25167447|1
nuclear receptor subfamily 2 group F me
cg04943986|1
cg08370996|1
U deubiquitinase with linear linkage spe
cg01794265|1
cg23639989|1
PBX homeobox 4
cg19996355|1
cg23158811|1
post-GPI attachment to proteins
cg02940164|1
cg21091629|1
protocadherin gamma subfamily B,
cg00911351|1
cg26282384|1
RAB32, member RAS oncogene far
cg14889768|1
cg20098887|1
ribosomal protein L31
cg19032144|1
cg22809047|1
secretagogin, EF-hand calcium binding
cg06493994|1
cg16954341|1
serine protease 12
cg07625840|1
cg21208104|1
Beta value
Age

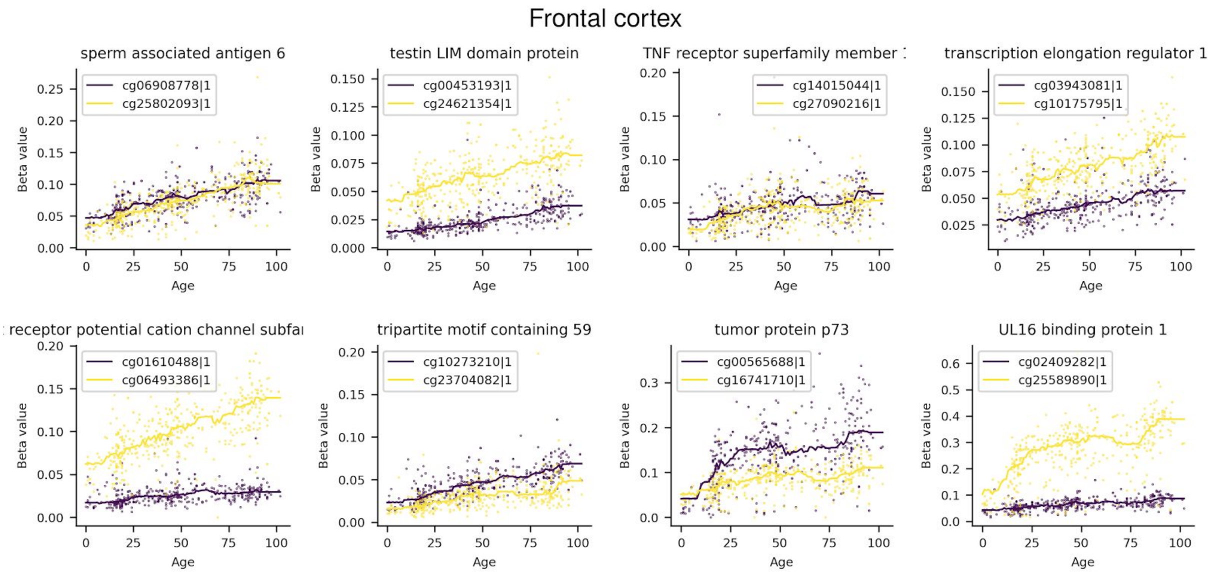


**Figure S4.** Genes for the Frontal cortex that are regulated by at least 2 significant markers. The marker identifier and the cluster label are separated by "|". All belong to cluster 1.

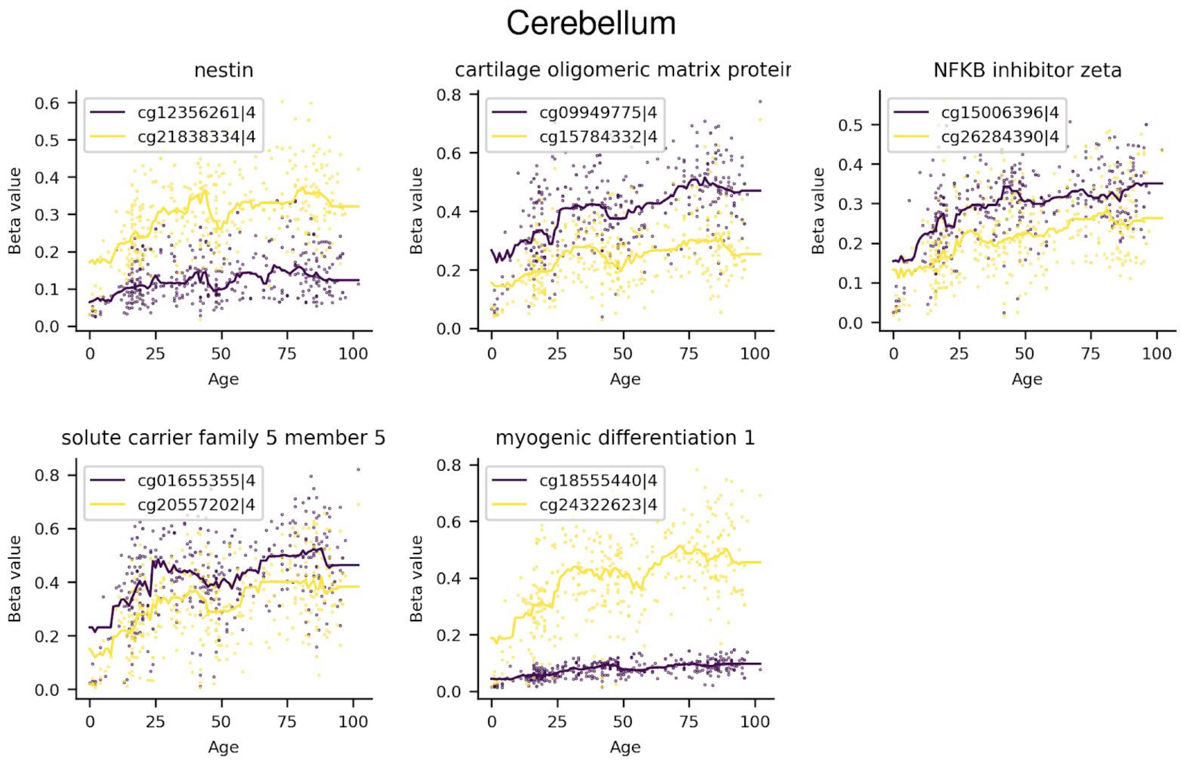


**Figure S5.** Genes for the Cerebellum that are regulated by at least 2 significant markers. The marker identifier and the cluster label are separated by "|". All belong to cluster 4.

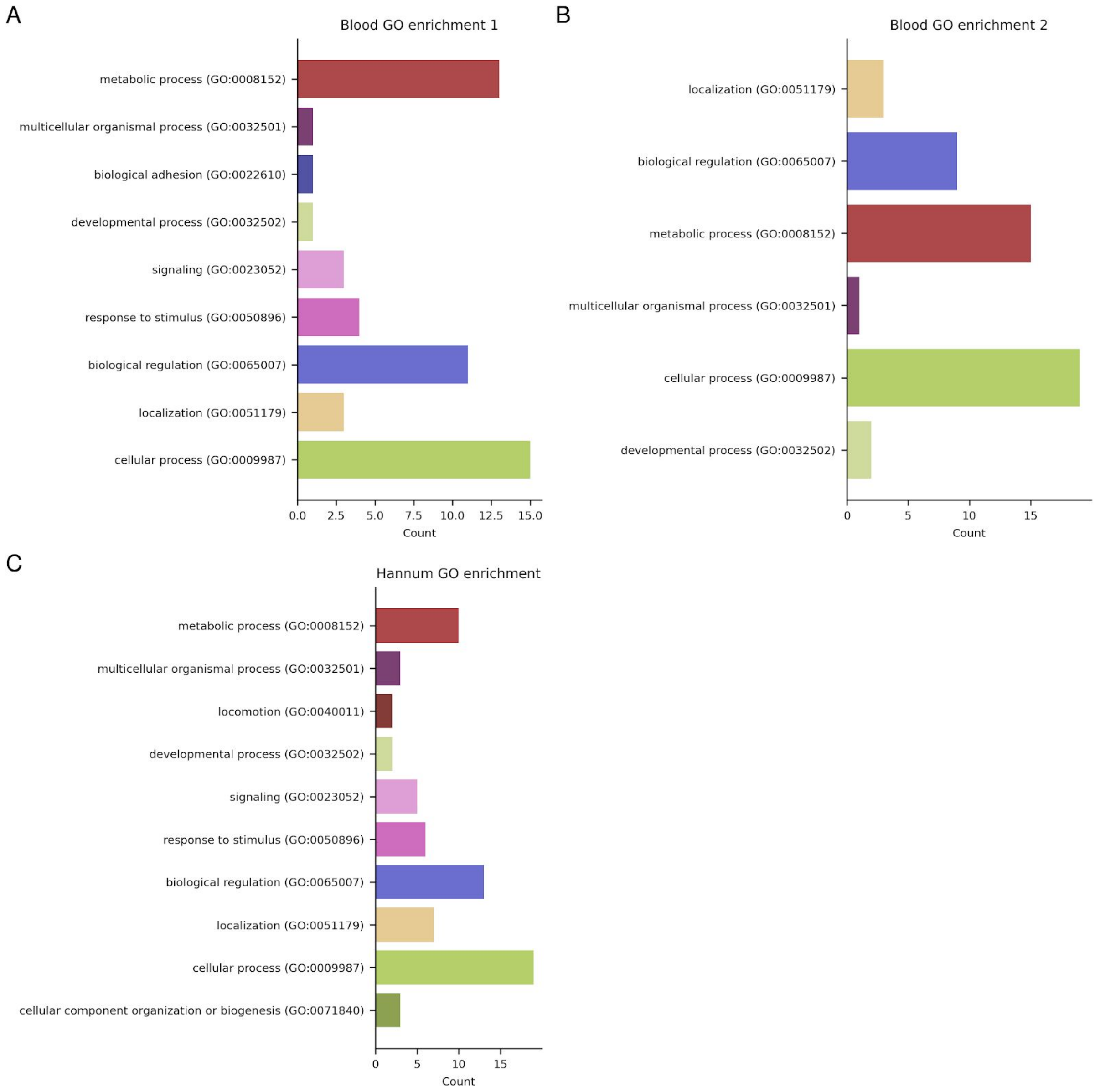


**Figure S6** Gene Ontology enrichment for the genes related to the blood cluster 1(**A**) and 2 (**B**) and Hannum (**C**) markers.

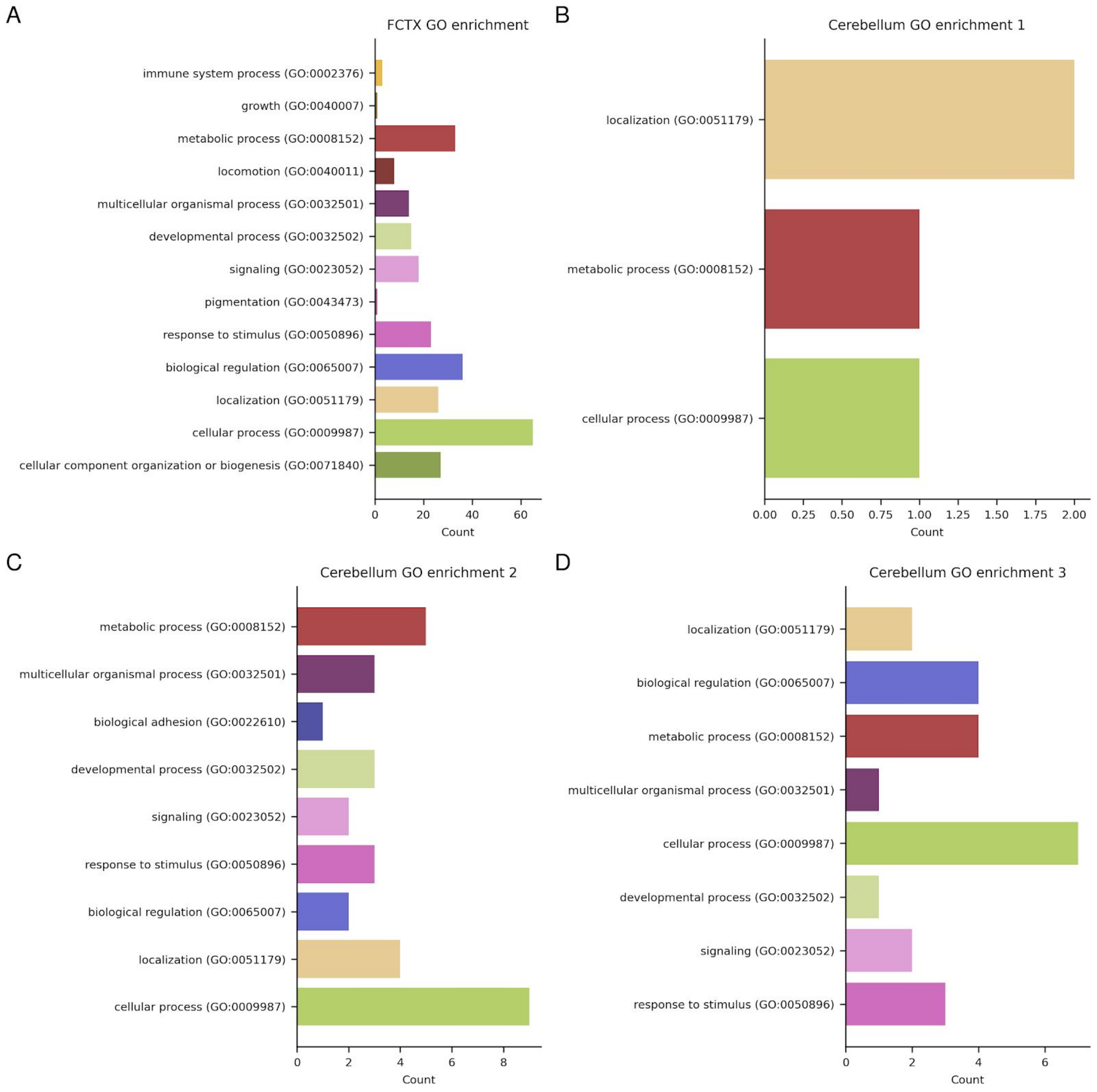


**Figure S7** Gene Ontology enrichment for the genes related to the frontal cortex (**A**) and cerebellum clusters 1-3 (**B-D**) markers.

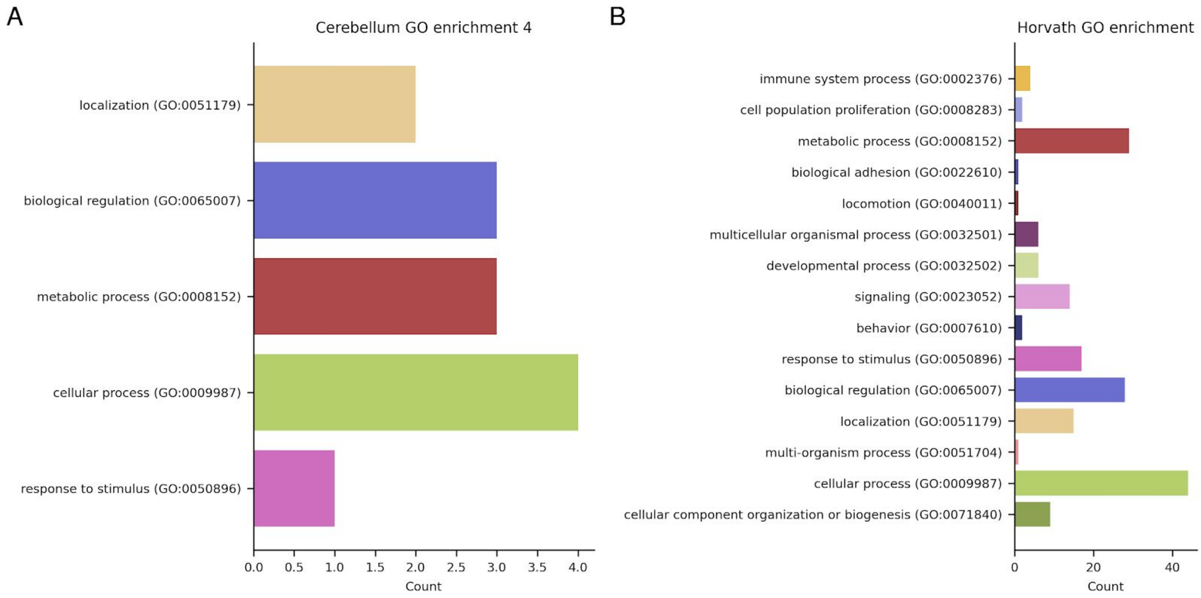


**Figure S8.** Gene Ontology enrichment for the genes related to cerebellum cluster 4 and the Horvath markers.
